# Granulin Loss of Function in Human Mature Brain Organoids Implicates Astrocytes in TDP-43 Pathology

**DOI:** 10.1101/2022.10.24.513566

**Authors:** Martina de Majo, Mark Koontz, Elise Marsan, Nir Salinas, Arren Ramsey, Yien-Ming Kuo, Kyounghee Seo, Huinan Li, Nina M Dräger, Kun Leng, Santiago L Gonzales, Michael Kurnellas, Yuichiro Miyaoka, Joseph R Klim, Martin Kampmann, Michael E Ward, Eric J Huang, Erik M Ullian

## Abstract

Loss of function (LoF) of Tar-binding protein 43 (TDP-43) and mislocalization, together with TDP-43-positive and hyperphosphorylated inclusions, are found in postmortem tissue of amyotrophic lateral sclerosis (ALS) and frontotemporal dementia (FTD) patients, including those carrying LoF variants in the progranulin gene (*GRN*). Modelling TDP-43 pathology has been challenging *in vivo* and *in vitro*. We present a 3D-induced pluripotent stem cell (iPSC)-derived paradigm - mature brain organoids (mbOrg) - composed of cortical-like-astrocytes (iA) and neurons (iN). When devoid of *GRN*, mbOrgs spontaneously recapitulate TDP-43 mislocalization, hyperphosphorylation and LoF phenotypes. Mixing-and-matching genotypes in mbOrgs showed that *GRN*^−/−^ iA are drivers for TDP-43 pathology. Finally, we rescued TDP-43 LoF by adding exogenous progranulin, demonstrating a link between TDP-43 LoF and progranulin expression. In conclusion, we present an iPSC-derived platform that shows striking features of human TDP-43 proteinopathy and provides a tool for mechanistic modelling of TDP-43 pathology and patient-tailored therapeutic screening for FTD and ALS.

**Highlights:**

- *GRN*^−/−^ iPSC-derived 3D paradigm (mbOrg) composed of mature cortical-like astrocytes and neurons recapitulates features of TDP-43 proteinopathy
- *GRN*^−/−^ cortical-like astrocytes are necessary and sufficient for the development of the TDP-43 loss of function phenotype in mbOrg.
- A TDP-43 phenotype can be rescued in neurons by treating neuron and astrocyte co-cultures with progranulin full length protein.

**eTOC blurb:** In this article, de Majo and colleagues present a novel 3D iPSC-derived model to study neurodegenerative disorders such as ALS and FTD. When devoid of *GRN* expression, these cultures present features of ALS and FTD associated pathology hardly ever observed *in vitro*. These phenotypes are shown to be primarily driven by diseased astrocytes and can be rescued by progranulin supplementation.

## Introduction

Frontotemporal dementia (FTD) and amyotrophic lateral sclerosis (ALS) are two fatal neurodegenerative diseases. ALS is estimated to affect 2.1 cases per 100000 people per year (Chiò et al., 2013) while FTD is the second most common cause of dementia for people under 65 (Knopman & Roberts, 2011). ALS and FTD are now thought to be different manifestations of the same disease spectrum with ALS primarily affecting the motor system and FTD presenting with a variety of symptoms affecting behavioral, executive, language, and motor functions. The two clinical entities can also occur in the same patients, with ~10-15% of ALS patients diagnosed with FTD features (FTD-ALS) and ~50% ALS patients developing some cognitive impairment (Burrell et al., 2016). TAR DNA Binding Protein (*TARDBP*) gene encodes for the TDP-43 protein, which plays a pivotal role in these two devastating neurodegenerative disorders. TDP-43 was also recently implicated in limbic-predominant age-related TDP-43 encephalopathy (LATE) (Nelson et al., 2022). Although TDP-43 is an RNA/DNA binding protein that physiologically resides in the nucleus, hyperphosphorylated extranuclear inclusions are found in neuronal cells of ~45% FTD and ~97% of ALS patients (Ling et al., 2013; Neumann et al., 2006). Loss of nuclear TDP-43 results in defective splicing of several transcripts among which the most thoroughly described is the cryptic splicing of Stathmin 2 (*STMN2*), a neuron-specific gene important for neuronal survival (Akiyama et al., 2022; Klim et al., 2019; Melamed et al., 2019; Prudencio et al., 2020).

Roughly 5-10% of all FTD patients harbor mutations in the granulin (*GRN*) gene. (Baker et al., 2006; Cruts et al., 2006; Seelaar et al., 2008). *GRN* transcripts are composed of 13 exons and encode for the progranulin (PGRN) protein, a highly conserved ~80 kDa protein involved in lysosomal function, neuronal survival, and inflammation (Rhinn et al., 2022). The vast majority of the *GRN* mutations are dominant heterozygous loss of function variants that lead to haploinsufficiency with consequent lower expression of PGRN. These patients have been reported to harbor TDP-43 proteinopathy in their frontal and/or temporal lobe at postmortem examination (Baker et al., 2006; Mackenzie, 2007). The *GRN* gene has been further linked to Alzheimer’s disease and LATE, suggesting this gene plays roles in multiple neurodegenerative diseases (Bellenguez et al., 2020; Nelson et al., 2019; Viswanathan et al., 2009)

To date, modelling TDP-43 proteinopathy *in vitro* or *in vivo* has been a challenge, with most *in vitro* models applying exogenous stress to replicate pathology akin to what is observed in postmortem ALS/FTD central nervous system (CNS) tissue. Within these paradigms, induced pluripotent stem cells (iPSCs) hold great promise as they allow for patient- and tissue-specific human-derived *in vitro* models, which can be used for therapeutic screening and phenotype testing (Palomo et al., 2019). As an example, recent work has established the utility of iPSC models for the study of *GRN* loss of function (LoF) (Lee & Huang, 2017; Rosen et al., 2011). *GRN* depletion in human iPSC-derived neurons causes cell autonomous changes in signaling (Almeida et al., 2012) and cellular stress (Rosen et al., 2011). Interestingly, these *in vitro* studies did not replicate the characteristic TDP-43 pathology observed in patients (Ahmed et al., 2010; Ward et al., 2014). One possible explanation for this is the fact that *GRN* is also expressed in glia, such as microglia and astrocytes, raising the likelihood that *GRN* loss of function causes pathology in a non-cell autonomous manner (Kelley et al., 2018; Y. Zhang et al., 2014, 2016). Recent studies have further implicated glial cells, including astrocytes and microglia, in neurodegenerative disease progression (Desai et al., 2010; Hansen et al., 2018; Verkhratsky et al., 2010). Indeed, several lines of evidence from *GRN* knockout mouse models indicate PGRN deficiency induces glial complement activation and non-autonomous microglia-mediated synaptic pruning that subsequently leads to neurodegeneration (Lui et al., 2016). The role of astrocytes, however, has been less thoroughly characterized and important questions remain such as whether disease-associated astrocytes are capable of inducing TDP-43 pathology and whether human models of FTD can show pathology similar to that found in postmortem human brain.

To address this challenge, we applied our previously developed iPSC-derived 3D co-culture model (Krencik et al., 2017; Liu et al., 2020) composed of mature cortical-like neurons and astrocytes, assembled in precise ratios and numbers, to study *GRN* LoF in FTD. When devoid of granulin expression (*GRN*^−/−^), our model develops features of TDP-43 pathology, including cryptic *STMN2* (CrSTMN2) splicing, and extranuclear and hyperphosphorylated TDP-43 inclusions. This study presents the first *in vitro* model showing robust evidence of numerous FTD/ALS pathology markers spontaneously developing, overcoming the need of exogeneous chemical-induced stress or overexpression. Furthermore, we obtained partial phenotype rescue when *GRN*^−/−^ cells were treated with exogenous full length PGRN, demonstrating that the development of TDP-43 pathology is dependent on PGRN expression. We believe this model could provide insight into cell biological mechanisms leading to TDP-43 pathology and offer a platform for patient tailored phenotype and therapeutic screening for FTD and ALS patients with suspected or confirmed TDP-43 proteinopathy.

## Results

To investigate the role of astrocytes in *GRN* LoF, we modified a previously described protocol to study glial-neuronal interactions (Krencik et al., 2017; **Fig 1 A**). This approach entails generating neurogenin 2 (NGN2) inducible cortical-like neurons (iNeurons, iN), which readily form synapses (Fernandopulle et al., 2018), and mature cortical-like astrocytes (iAstrocytes, iA) (Krencik et al., 2015; Krencik & Zhang, 2011), and assembling them into 3D organoid-like structures at defined numbers and ratios of neurons and astrocytes (termed Mature Brain Organoids [mbOrgs])(**Fig 1 B**). This approach allows us to better model the ratio of astrocytes to neurons thought to comprise the human cortex as well as mix and match neurons and astrocytes derived from either isogenic wildtype (*GRN*^+/+^) iPSCs or isogenic *GRN* knockout (*GRN*^−/−^) iPSCs (sFig2G). Importantly, *GRN*^−/−^ mbOrgs show a complete loss of PGRN (s**Fig 1 B,E**). Using this approach, we interrogated the pathological phenotypes of *GRN*^+/+^ and *GRN*^−/−^ mbOrgs. Using transmission light microscopy and confocal imaging, we showed that mbOrgs formed from *GRN*^+/+^ or *GRN*^−/−^ iPSCs both developed into uniform spheres containing a readily detectible array of astrocytes and neurons (**Fig 1, B, C**). We first looked at standard markers characteristic of iPSC-derived neurons and astrocytes, by immunostaining. Aquaporin 4 (*AQP4*), an astrocyte specific gene well expressed by astrocytes (Krencik et al., 2015), was strongly and uniformly expressed in both *GRN*^+/+^ and *GRN*^−/−^ mbOrgs (**Fig 3A**). Similarly, we found a strong and widespread expression of neuronal microtubule associated protein 2 (MAP2) expression in both mbOrgs (**Fig 3B**). Both results indicate overall stable expression of these astrocyte and neuronal markers in *GRN*^+/+^ as well as *GRN*^−/−^ mbOrgs.

**Figure 1:**
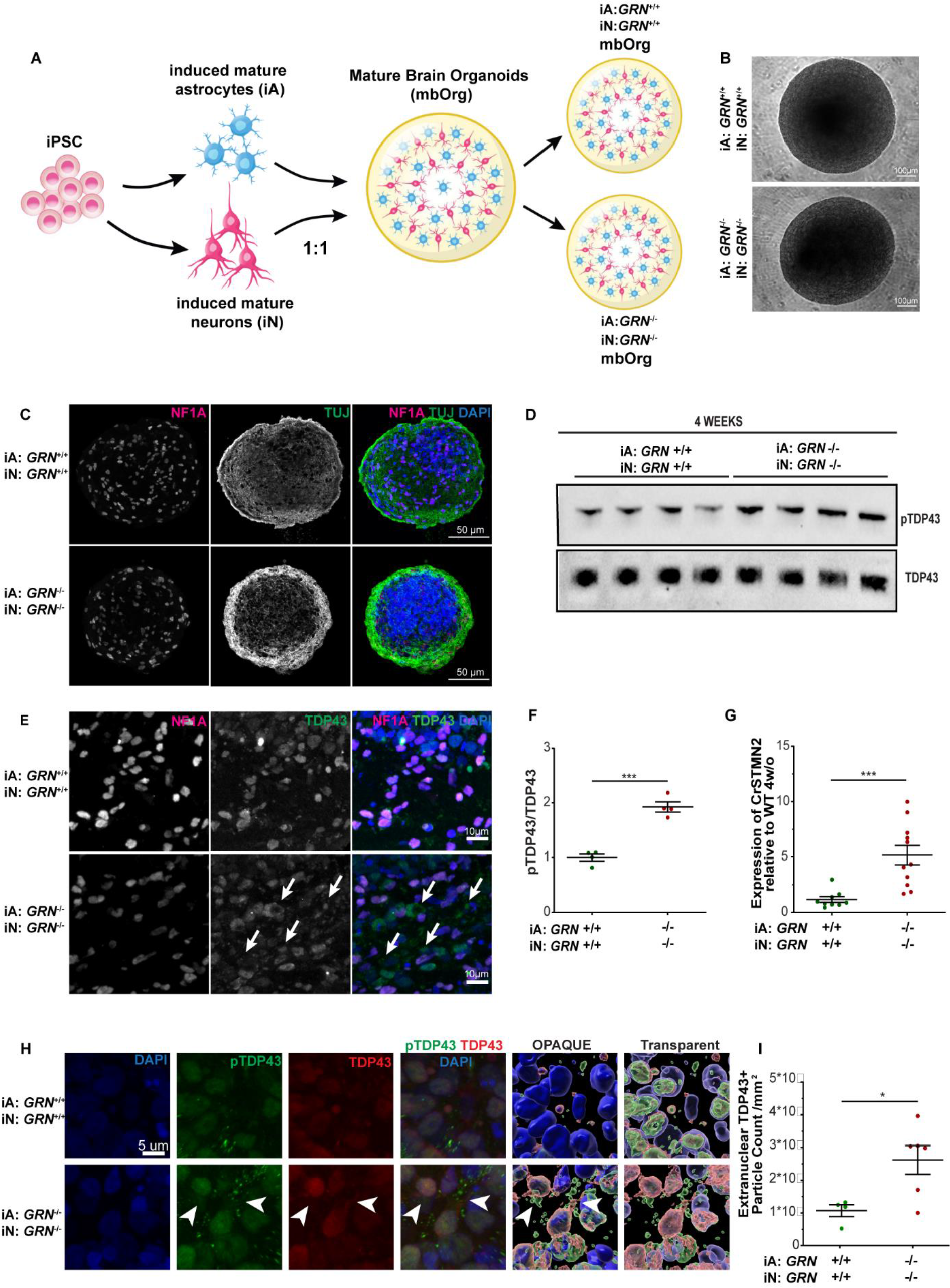
GRN^−/−^ Mature Brain Organoids (mbOrg) show features of human TDP-43 proteinopathy after four weeks in culture. A) Diagram showing the process by which mbOrg are differentiated and assembled. Briefly, iPSCs are differentiated in mature astrocytes and cortical-like neurons and assembled in a 1:1 ratio in 3D cocultures called mbOrg. The mbOrg are grown for four weeks and then analyzed. B) Brightfield image of two organoids (*GRN^+/+^*, *GRN^−/−^*) kept in culture for four weeks (scale bar 100μm). C) Confocal images of organoid slices (*GRN^+/+^* or *GRN^−/−^*) showing expression of NF1A, TUJ and DAPI after four weeks in culture (scale bar 50 μm). D) Western blot of mbOrg whole lysate showing higher expression on pTDP-43 in *GRN^−/−^* compared to *GRN^+/+^* mbOrg when normalized to total TDP-43. F) Western blot quantification showing significantly higher expression on pTDP-43 in *GRN^−/−^* compared to *GRN^+/+^* mbOrg when normalized to total TDP-43 (n=4, unpaired t test, two tailed, p<0.001). E) Confocal images of organoid slices (*GRN^+/+^* or *GRN^−/−^*) showing expression of NFIA, TDP-43 and DAPI after four weeks in culture (scale bar 10 μm). G) Quantification of cryptic STMN2 (CrSTMN2) expression using qPCR showing significantly CrSTMN2 higher expression in *GRN^−/−^* compared to *GRN^+/+^* mbOrg (n=11±, unpaired t test, two tailed, p=<0.001). H) IMARIS 3D reconstruction of TDP-43 and pTDP-43 staining in four week old mbOrg (scale bar 5 μm). I) quantification of extranuclear TDP-43 particle count per mm^2^ showing it to be significantly higher in *GRN^−/−^* compared to *GRN^+/+^* mbOrg. Each dot represents one independent mbOrg (*GRN^+/+^* n=4, *GRN^−/−^* n=6, unpaired t test, two tailed, p<0.05). For all graphs data are represented as mean ± SEM.

*GRN* associated FTD TDP-43 proteinopathy is characterized by increased levels of phospho-TDP-43 (pTDP-43) and abnormal TDP-43 cytoplasmic accumulation, seen in postmortem CNS tissue (Baker et al., 2006; Mackenzie, 2007). To assess the TDP-43 pathology in our model, we measured the levels of both pTDP-43, using a ser409/410 specific pTDP-43 antibody, and total TDP-43 by performing Western Blot in *GRN*^+/+^ and *GRN*^−/−^ mbOrgs lysates after four weeks of culture (**Fig 1D**). Quantification of the pTDP-43 to TDP-43 ratio revealed a clear increase in the phosphorylated form of TDP-43 in *GRN^−/−^* mbOrgs (**Fig 1 F**); similar to what has been described in postmortem tissue (Baker et al., 2006; Mackenzie, 2007). We next performed immunostaining for TDP-43 in sections of four weeks old mbOrgs and found clear evidence of extra-nuclear localization of TDP-43 in *GRN*^−/−^ mbOrgs but not in *GRN*^+/+^ mbOrgs, where TDP-43 was mostly colocalized with nuclei (**Fig 1E**). IMARIS reconstruction and quantification of confocal images from TDP-43 staining demonstrated an increase in extranuclear TDP-43 particle count in *GRN*^−/−^ mbOrgs (**Fig 1H, 1I**). Recent work has shown that a key function of TDP-43 in healthy cells is mRNA splicing repression in the nucleus, whereas in disease TDP-43 nuclear depletion results in a number of mis-spliced transcripts (Brown et al., 2022). Among these, *STMN2* is the most thoroughly studied; mis-splicing of *STMN2* transcripts has been considered a robust indicator of TDP-43 pathology and correlates with the level of pTDP-43 (Klim et al., 2019; Melamed et al., 2019; Prudencio et al., 2020). Thus, to determine if cryptic *STMN2* transcripts (CrSTMN2) can be detected in our model system showing TDP-43 mislocalization, we adapted a method (Klim et al., 2019) to develop a sensitive qPCR-based assay and found a highly significant, approximately 4-fold increase in CrSTMN2 in the *GRN^−/−^* vs. *GRN^+/+^* mbOrgs (**Fig 1G**). We next asked if these unique features of human neurodegenerative disease progress over time in *GRN*^−/−^ mbOrgs, as would be expected if these features represent cellular mechanisms that could be relevant to human disease progression. To test this, we looked at an earlier two-week time point and assessed the presence of these same features that are robustly present at four weeks. We found that, while there was a trend, variable results caused these features to have not yet reached statistical significance at the earlier time point, consistent with a progression of relevant cellular signaling driving these features (**sFig 1A,C,D**). Taken together, these data demonstrate a remarkable degree of human specific FTD pathological phenotypes recapitulated in the iPSC-derived mbOrgs model and are, to the best of our knowledge, the first demonstration of multiple TDP-43 associated pathological phenotypes shown in an unperturbed *in vitro* model system.

These results show that *GRN* loss of function in both iPSC-derived neurons and astrocytes in our 3D platform display a remarkable array of phenotypes relevant to FTD-TDP. We next wanted to investigate if *GRN* loss of function is required in both cell types or if we can detect evidence of pathology when either neurons or astrocytes are *GRN*^−/−^. To investigate this, we took advantage of the assembled nature of the mbOrgs and made heterotypic cultures containing all possible combinations of *GRN*^−/−^ or *GRN^+/+^* neurons + *GRN^−/−^* or *GRN^+/+^* astrocytes (either *GRN*^−/−^ neurons + *GRN*^+/+^ astrocytes or *GRN*^+/+^ neurons *+ GRN^−/−^* astrocytes, along with the control both-cell-type *GRN*^+/+^ and both-cell-type *GRN*^−/−^ mbOrgs). Immunostaining for TDP-43 showed expected pathology in the both-cell-type *GRN*^−/−^ mbOrgs. We also found TDP-43 pathology in the *GRN*^+/+^ neurons + *GRN*^−/−^ astrocytes mbOrgs (**Fig 2A**). To further quantitatively examine the heterotypic cultures, we assessed CrSTMN2. As expected, quantification of CrSTMN2 showed the most severe phenotype in both-cell-type *GRN*^−/−^ mbOrgs. Surprisingly, we found a robust CrSTMN2 increase in *GRN*^+/+^ neurons + *GRN*^−/−^ astrocytes mbOrgs (**Fig 2B**) but not in *GRN*^−/−^ neurons + *GRN*^+/+^ astrocytes mbOrgs, confirming what was observed by immunocytochemistry. These results demonstrate that *GRN*^−/−^ astrocytes are sufficient to induce robust TDP-43 and CrSTMN2 phenotypes in mbOrgs, even in presence of *GRN*^+/+^ neurons and indicate that the diseased human astrocytes can drive neurodegenerative phenotypes in healthy human neurons.

**Figure 2:**
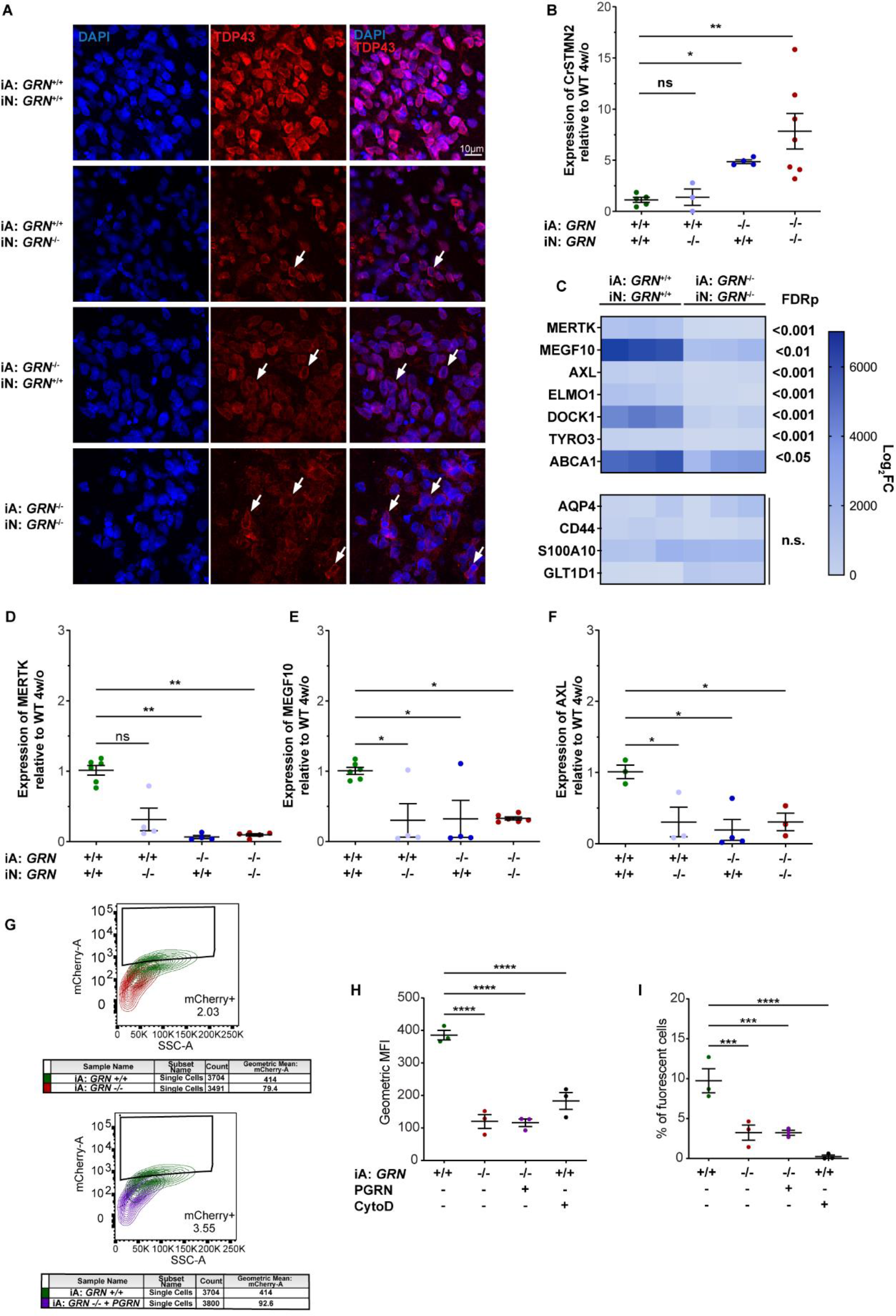
GRN^−/−^ mature astrocytes drive STMN2 mis-splicing in mbOrg and show evidence of defective phagocytosis. A) Qualitative immunohistochemistry of mbOrgs slices (*GRN* iA: +/+ iN: +/+, *GRN* iA: +/+ iN: −/−, *GRN* iA: −/− iN: +/+, *GRN* iA: −/− iN: −/−) showing expression of TDP-43 DAPI after four weeks in culture (scale bar 5 μm). B) Quantification of cryptic STMN2 (CrSTMN2) expression using qPCR showing increasingly significant CrSTMN2 higher expression in *GRN* iA: −/− iN: +/+ and *GRN* iA: −/− iN: −/− compared to *GRN* iA: +/+ iN: +/+ and *GRN* iA: +/+ iN: −/− mbOrg (n=4, one way Anova followed by multiple comparison, *p<0.05 **p<0.005). C) At the top expression heat map showing bulk RNA sequencing values of 7 genes linked to phagocytosis in four week old mbOrgs. All the genes listed here are differentially expressed in the *GRN^+/+^* samples vs *GRN^−/−^* samples. Specifically, all the genes listed here less expressed in the four week old *GRN^−/−^* compared to the *GRN^+/+^* mbOrgs (FDR<0.05, n=3). At the bottom expression heat map of astrocyte markers that are not differentially expressed in the four week old *GRN^−/−^* compared to the *GRN^+/+^* mbOrg. Fold Change (FC). D,E,F) qPCR analysis of three (MERTK, MEGF10 and AXL) of the seven phagocytosis markers analyzed with RNAseq in all four conditions: *GRN* iA: +/+ iN: +/+, *GRN* iA: +/+ iN: −/−, *GRN* iA: −/− iN: +/+, *GRN* iA: −/− iN: −/− (n=4, one way Anova followed by multiple comparison, *p<0.05 **p<0.005). G) Graphic representation of the clustering analysis depicting the changes of phagocytic activity in the different conditions (iA *GRN^+/+^* in green, iA *GRN^−/−^* in red and iA *GRN^−/−^* + PGRN in purple). Cells were treated with mCherry labelled rat synaptosome and analyzed at the cytofluorimeter. H, I) Quantification of FACS analysis showing geometric mean fluorescence intensity (MFI) and percentage of fluorescent cells phagocyted by the mature astrocytes. iA *GRN^−/−^* and iA *GRN^−/−^* + PGRN phagocyte significantly lower amount of mCherry labelled synaptosome according to both parameters compared to iA *GRN^+/+^*. Negative control showed in black (iA *GRN^+/+^* treated with cytoD). For all graphs data are represented as mean ± SEM.

To further characterize the effect of *GRN* loss of function on iAstrocytes we performed bulk RNA sequencing (**sFig2D-E**) to examinee gene expression in *GRN*^+/+^ and *GRN*^−/−^ mbOrgs after four weeks in culture. Interestingly, we found that a set of astrocyte genes known to be involved in phagocytosis (**Fig 2C upper panel**), but not other astrocyte specific marker genes, were down-regulated in the *GRN*^−/−^ mbOrgs at four weeks when compared to *GRN*^+/+^ mbOrgs (**Fig 2C lower panel**). Confirmation with qPCR, shows MER Proto-Oncogene, Tyrosine Kinase (MERTK), multiple EGF like domains 10 (MEGF10), and AXL receptor tyrosine kinase (AXL) down-regulation in *GRN*^−/−^ mbOrgs. Interestingly, all three phagocytosis-related genes were also significantly down-regulated in mbOrgs composed of *GRN*^+/+^ neurons + *GRN*^−/−^ astrocytes. For the *GRN*^−/−^ neurons + *GRN*^+/+^ astrocytes mbOrgs both MEGF10 and AXL were significantly down-regulated and there was a non-significant trend for down regulation for MERTK (**Fig 2D,E,F**). These results suggest that *GRN* loss of function in astrocytes and/or neurons can lead to a down regulation of phagocytosis-related astrocytes genes. To directly assess the functional consequences of *GRN* loss in astrocytes, we performed a synaptosome phagocytosis assay in *GRN*^+/+^ or *GRN*^−/−^ astrocytes 2D cultures. The results point to a profound deficit in synaptosome phagocytosis in the *GRN*^−/−^ astrocytes (**Fig 2G,H,I**). We attempted to rescue this deficit with the addition of recombinant progranulin (PGRN), however, the tested conditions were not sufficient to revert the phagocytosis deficit (**Fig 2G,H,I**). A similar treatment with PGRN in 2D astro-neuronal cultures for up to four weeks failed to rescue the phagocytosis-related down-regulated genes MERTK and MEGF10 (**sFig1H and K**). Overall, these data show that *GRN* loss of function leads to changes in astrocyte phagocytosis and that the genes associated with astrocyte phagocytosis are regulated both cell autonomously and non-cell autonomously.

Previous work has uncovered a critical role for astrocyte phagocytosis in regulating synapses in various regions of the mouse CNS with specific involvement of MERTK and MEGF10 (Chung et al., 2013). We wondered whether the deficits we observed in *GRN*^−/−^ astrocyte phagocytosis may be associated with corresponding increases in synapses in our model. Therefore, we looked for any synaptic phenotypes associated with loss of phagocytosis in the mbOrgs. First, we immunostained mbOrgs for the presynaptic marker synaptophysin (SYP) and postsynaptic marker postsynaptic density protein 95 (PSD-95). Confocal microscopy analysis and IMARIS 3D reconstruction showed that in comparison to four-week-old *GRN^+^*^/+^ mbOrgs, *GRN*^−/−^ mbOrgs contain more SYP and PSD-95 puncta (**Fig 3C**). We next wanted to quantitatively assess the expression levels of synaptic markers and performed Western blot for both pre- and post-synaptic markers synapsin (SYN1) and PSD-95, respectively. The results revealed an increase in both pre- and post-synaptic protein in the *GRN*^−/−^ mbOrgs (**Fig 3D,E**). Thus, *GRN* loss is associated with effects on synapses consistent with previous studies (Petoukhov et al., 2013; Tapia et al., 2011; Uesaka et al., 2018; L. Wang et al., 2022).

**Figure 3:**
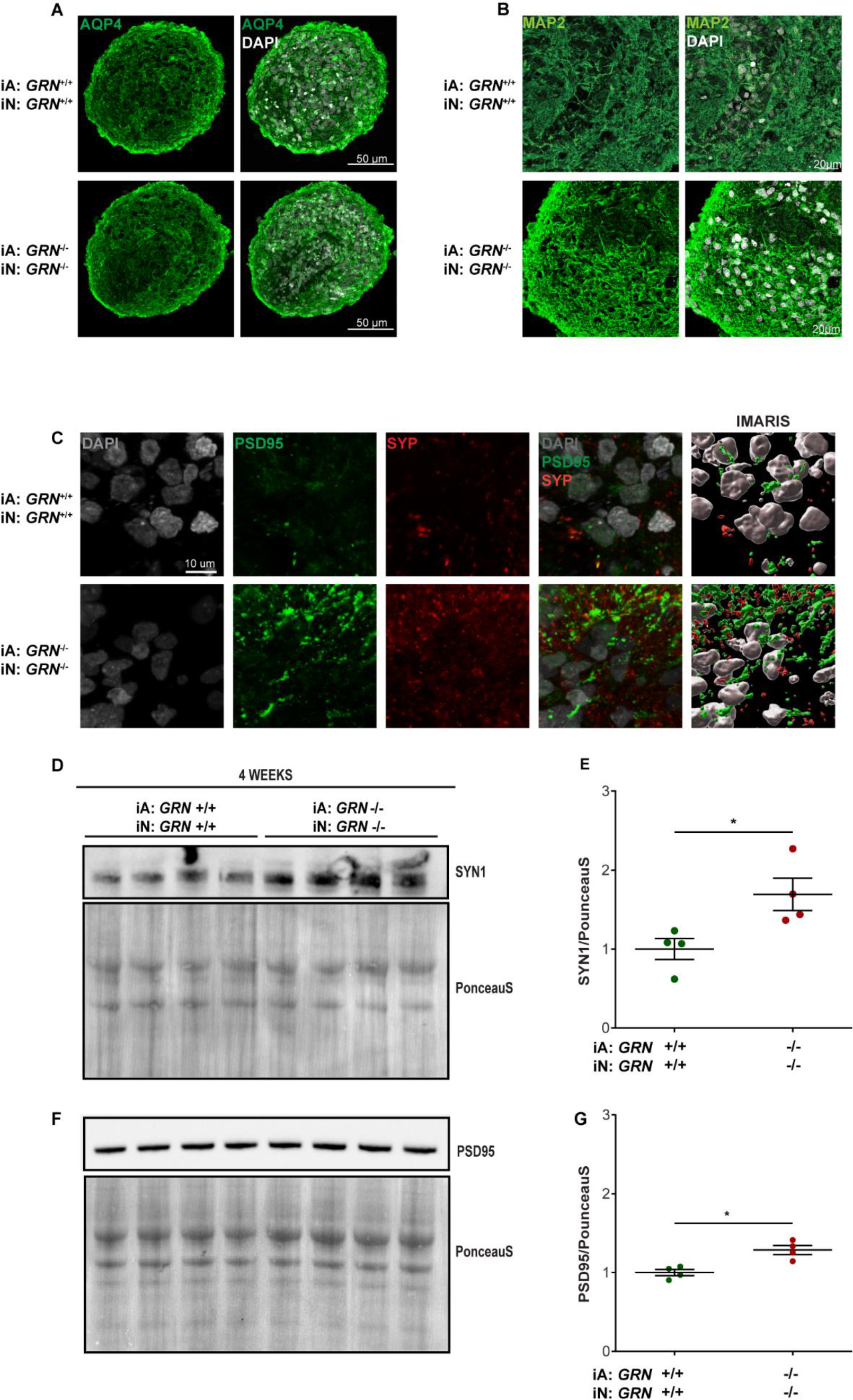
GRN^−/−^ mbOrg show higher synaptic density when compared to GRN^+/+^ mbOrg. A) Confocal images of organoid slices (*GRN^+/+^* or *GRN^−/−^*) showing evenly distributed expression of AQP4 and DAPI (scale bar 50 μm). B) Confocal images of organoid slices (*GRN^+/+^* or *GRN^−/−^*) showing evenly distributed expression of MAP2 and DAPI (scale bar 20 μm). C) IMARIS 3D reconstruction of PSD95 and Synaptophysin (SYP) staining in four week old mbOrg (scale bar 10 um) showing higher synaptic density in *GRN^−/−^* compared to *GRN^+/+^* mbOrg. D) Western blot of mbOrg whole lysate showing higher expression on Synapsin1 (SYN1) in *GRN^−/−^* compared to *GRN^+/+^* mbOrg when normalized to PonceauS staining and its quantification (E, n=4, unpaired t test, two tailed, p<0.05). F) Western blot of mbOrg whole lysate showing higher expression on PSD95 in *GRN^−/−^* compared to *GRN^+/+^* mbOrg when normalized to PonceauS staining and its quantification (G, n=4, unpaired t test, two tailed, p<0.05). For all graphs data are represented as mean ± SEM.

Overall, these findings indicate that *GRN*^−/−^ astrocytes are likely sufficient to induce TDP-43 pathology and a significant increase in CrSTMN2 in neurons in the 3D mbOrgs. We then decided to investigate if the same two findings are present in a 2D culture system. We performed the co-culture experiments as illustrated in **Figure 4A**. Both *GRN*^+/+^ and *GRN*^−/−^ co-cultures of astrocytes and neurons look uniformly healthy at four weeks (28 DIV) (**Fig 4B**). When immunostained for MAP2, both cultures showed robust and extensive dendritic arbors (**Fig 4B**). As previously reported, staining for TDP-43 and pTDP-43 failed to show an overt pathological TDP-43 associated signal. Furthermore, Western blot analysis for the ratio of pTDP-43 to total TDP-43 showed no significant increase at four weeks in *GRN^−/−^* 2D co-cultures. We then assessed the expression of CrSTMN2 in the 2D co-cultures. We found a significant increase in the expression of CrSTMN2 at four weeks in 2D co-cultures of *GRN*^−/−^ astrocytes with *GRN*^−/−^ neurons (**Fig 4G**). This result led us to investigate if this more-subtle TDP-43 associated phenotype might be reversible. Although phenotypes are not as severe in the 2D co-culture, they are amenable to exogenous compound rescue experiments. As proof of principle, we tried rescuing the CrSTMN2 phenotype by treating the cells with recombinant PGRN. We first determined what, if any, concentration of PGRN shows clear cellular uptake. We determined by immunostaining that 1μg/ml of recombinant PGRN fed every three days for 28 days leads to a robust rescue of PGRN deficiency in *GRN*^−/−^ neuronal and *GRN*^−/−^ astrocytes cell bodies (**Fig 4F**). We next examined CrSTMN2 in PGRN-treated cells *vs*. control and found a consistent and significant rescue of CrSTMN2 in 2D co-cultures of *GRN*^−/−^astrocytes and *GRN*^−/−^ neurons (**Fig 4G**). These are, to our knowledge, the first data demonstrating a rescue of *GRN* loss of function CrSTMN2 expression increases.

**Figure 4:**
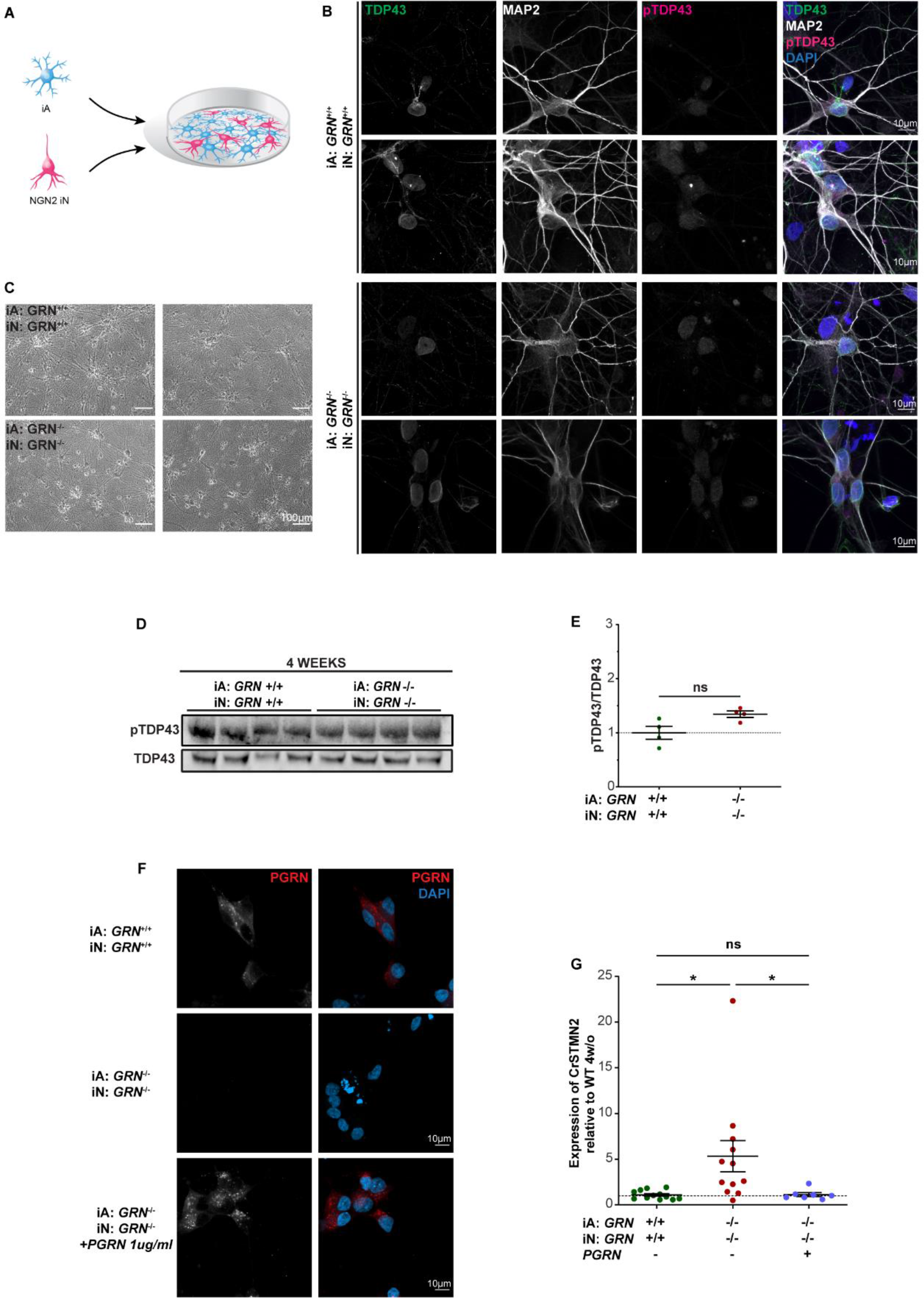
STMN2 mis-splicing can be rescued in 2D. A) Diagram showing the process by which 2D cultures are made. Briefly, iPSCs are differentiated in mature astrocytes and cortical-like neurons and assembled in a 1:1 ratio in 2D cocultures. The cocultures are grown for four weeks and then analyzed. B) Representative ICC images of 2D cultures stained for TDP-43, pTDP-43, MAP2 and DAPI (scale bar 10μm). C) Brightfield image of 2D cultures (*GRN^+/+^*, *GRN^−/−^*) at four week timepoint (scale bar 100μm). D) Western blot of 2D cocultures whole lysate of pTDP-43 and total TDP-43 in *GRN^+/+^* and *GRN^−/−^* 2D cocultures. E) western blot quantification showing similar expression on pTDP-43 in *GRN^−/−^* compared to *GRN^+/+^* 2D cocultures when normalized to total TDP-43 (n=4, unpaired t test, two tailed, p>0.05). F) Representative ICC images of 2D cultures stained for PGRN and DAPI showing positive staining in both *GRN^+/+^* and PGRN treated *GRN^−/−^* but no PGRN staining in *GRN^−/−^* cocultures (scale bar 10μm). G) Quantification of CrSTMN2 expression using qPCR showing significantly CrSTMN2 higher expression in *GRN^−/−^* compared to *GRN^+/+^* 2D cocultures. The difference is rescued when *GRN^−/−^* are treated with PGRN for four weeks (n=10, unpaired t test, one way Anova followed by multiple comparison, *p<0.05, ns p>0.05). For all graphs data are represented as mean ± SEM.

## Discussion

Here, we characterize an *in vitro* human iPSC-derived neuro-glia 3D model of TDP-43 proteinopathy in a context of PGRN deficiency. While iPSC induced neurons have provided numerous cell-autonomous biological insights into neurodegeneration, most models do not recapitulate multiple aspects of TDP-43 associated pathology. These phenotypic challenges have made therapeutic discovery difficult for TDP-43 proteinopathies (Buratti, 2020). Our focus on iPSC neuron-glial interactions have yielded a 3D paradigm spontaneously reproducing overt pathological features of TDP-43 proteinopathy. Interestingly, while TDP-43 proteinopathy was not noticeable in 2D co-culture, TDP-43 loss of function characterized by STMN2 mis-splicing was recapitulated in both 2D and 3D co-culture models.

Our approach is distinct in that we use a simple, reproducible, and straightforward engineered system to model spontaneous TDP-43 pathology. Indeed, our 3D iPSC induced cortical-like neuron and astrocyte co-culture with *GRN* loss strikingly and consistently recapitulated human specific TDP-43 cell pathology reported in human FTD patient brain. Importantly, these phenotypes are not seen in the same individually cultured iPSC induced neuron or astrocytes and are much milder in our 2D co-culture paradigm. Similarly, two recent studies highlighted evidence of mild TDP-43 pathology in 2D ALS-iPSC-derived neuron and 100-day-old *GRN* knock out neuronal cultures when compared to healthy controls, with the ALS study showing evidence of STMN2 mis-splicing (Bossolasco et al., 2022; Coyne et al., 2021). The reason for difference in phenotype severity between 2D and 3D co-cultures is not entirely clear, but we have previously found that astrocytes in 3D culture are significantly more complex and resemble the highly complex morphology of human astrocytes *in vivo* (Krencik et al., 2017). Furthermore, these previously published results suggest that control iPSC-derived human astrocytes in 3D are not highly reactive by four weeks *in vitro* (Krencik et al., 2015) potentially providing a healthy baseline to compare disease-causing mutations to.

The data presented here provide strong evidence that *GRN^−/^*^−^ astrocytes drive STMN2 mis-splicing in both *GRN^−/−^* and *GRN^+/+^* neurons in mbOrg. Indeed, the 3D mbOrgs are advantageous in allowing easy mix-and-matching of cellular genotypes with cells used to assemble the cultures. Thus, our ability to generate mixed *GRN^+/+^* neurons + *GRN^−/−^* astrocytes showing nearly as strong crSTMN2 and TDP-43 phenotypes of full *GRN^−/−^* neuron and astrocyte cultures indicate that *GRN* loss of function in human astrocytes can lead to TDP-43 loss of function in neurons. This finding is mechanistically important and adds to a growing body of literature suggesting that disease associated astrocytes and more generally glia can drive cell death and neuronal dysfunction (Huang et al., 2022; Leng et al., 2021; Liddelow et al., 2017; Taha et al., 2022; Zhang et al., 2020)

Our investigation of phagocytic activity assay in *GRN*^−^/^−^ iAstrocytes demonstrated significant deficit when compared to *GRN^+/+^* iAstrocytes. Phagocytic changes have been reported in GRN^−/−^ microglia (Guan et al., 2020; Lui et al., 2016) and in microglia differentiated from ALS peripheral blood mononuclear cells (PBMCs) when compared to control PBMCs-derived microglia (Quek et al., 2022). It has also been shown that diseased induced astrocytes lose the ability to engulf synapses and show reduced expression of phagocytosis receptor *MEGF10* and *MERTK* (Liddelow et al., 2017). We show that *GRN*^−/−^ iAstrocytes when cultured in 3D mbOrgs for four weeks also display differential expression of phagocytosis markers when compared to *GRN^+/+^* iAstrocytes. Significantly lower expression of *MERTK*, *MEGF10* and *AXL* was confirmed by qPCR in *GRN*^−/−^ iAstrocytes in both 3D and 2D co-cultures and was not rescued by PGRN treatment. *MEGF10* and *MERTK* have been specifically implicated in synaptic pruning and maturation in both in the developing and adult brain (Chung et al., 2013; Lee & Chung, 2019), their downregulation could explain the significantly higher expression of pre- and post-synaptic markers observed in our 3D *GRN*^−/−^ mbOrgs when comparted to *GRN^+/+^*. Overall, this model recapitulates key disease astrocyte-like features observed in neurodegenerative disorders, and it is, therefore, not surprising that *GRN*^−/−^ iA are able to drive the disease phenotype in mbOrgs even when co-cultured with healthy neurons.

As noted, our use of the mbOrgs revealed a compelling set of FTD-related phenotypes associated with TDP-43 loss of function. This finding gave us confidence that the significant crSTMN2 increase in the 2D co-cultures of neurons and astrocytes was caused by the lack of expression of *GRN*. Thus, we tested the ability of recombinant PGRN to rescue this deficit in the context of *GRN*^−/−^ neurons and astrocytes. Our results show a striking ability to rescue crSTMN2 increases in the 2D culture, consistent with this observation being a direct result of loss of *GRN* and subsequent loss of PGRN function. It is important to note that although treatment with PGRN can rescue crSTMN2 in the 2D co-culture, we did not find that treatment with PGRN was sufficient to rescue the observed phagocytosis deficits in the *GRN*^−/−^ iAstrocytes. This might be because these astrocytes had been matured for 6-9 months prior to analysis, so the changes associated with loss of *GRN* may not be acutely rescuable. We believe that the multifunctional PGRN protein is likely to affect numerous aspects of cellular function; some acutely and some chronically; thus, some deficits may be readily rescued by replacing PGRN and some may not. This is consistent with the recent finding that acute treatment with an engineered PGRN protein rescued some phenotypes of *GRN* deficient microglia but not all phenotypes associated with *GRN* loss (Logan et al., 2021).

Like all model systems, iPSC-derived models have limitations. In particular, they do not fully recapitulate all features of the complex human brain and they currently do not incorporate microglia or vascular networks, although this may be possible in the future (Blanchard et al., 2021; Kumar et al., 2017; Mantle & Lee, 2018). This paradigm (both in 2D and 3D) is, however, amenable to the addition of any further cell types to make it an even more complete modelling tool. Despite these limitations, this approach provides a simple straightforward method to model neurodegeneration that can be readily incorporated into laboratories and expanded to study basic cellular interactions and mechanisms of disease, revealing cell autonomous and non-autonomous roles for disease genes in astrocytes and neurons.

## Experimental procedures

### Human iPSC stem cell lines

Isogenic human iPSC line WTC11 and GRN−/− iPSC line were generated by Dr. Bruce R. Conklin as previously described (Miyaoka et al., 2014). *GRN*^−/−^ iPSC and *GRN*^−/−^ NGN2 iPSC were engineered (**sFig2F**) and provided by Dr. Michael E. Ward (NIH) as previously described (C. Wang et al., 2017). iPSCs were cultured and maintained in Essential 8 Medium (Gibco, A1517001) on 6 well cell culture plates (Olympus, 25-105) coated with Vitronectin (Gibco, A14700) in DPBS. iPSCs were dissociated and passaged using EDTA (Invitrogen, AM9260G) in DPBS.

### Cortical-like neuronal induction

Cortical-like iNeurons (iN) were generated as previously described (Fernandopulle et al., 2018). Briefly, iPSCs (WTC11) were expanded, dissociated, and replated on 10μg/ml Matrigel (Corning, 354234) coated plates. Cells were grown in specialized iNeuron induction media (iNIM) (DMEM-F12 + Glutamax (Gibco, 10565-018), N-2 supplement (Gibco, 17502-048), MEM-NEAA (Gibco, 11140-050) containing doxycycline (Sigma, D3072) for ~72 hrs, with media changed every ~24hrs. Cells were then dissociated using Accutase (Gibco, A1110501) and frozen in media + 10% DMSO (Sigma, D8418) at high density to maximize cell viability.

### Cortical-like astrocyte induction

Cortical-like iAstrocytes (iA) were generated as previously described (Krencik & Zhang, 2011). Human induced pluripotent stem cells (WTC11) were grown on vitronectin coated tissue culture plates using Essential 8 media. On day 0 of differentiation, iPSCs were dissociated into small aggregates averaging 50μm in diameter and transferred untreated tissue culture flasks with Neurosphere Induction Media (NSIM) (DMEM-F12/Neurobasal-A at 1:1 (Gibco, 10565-018: 108888-022)), N2 Supplement (Gibco, 17502-048), B27 -Vit.A Supplement (Gibco, 12587-010), MEM-NEAA (Gibco, 11140-050) plus SMAD inhibitors SB431542 (Stemcell Tech, 72234) and DMH1 (Tocris, 73634). Neurosphere induction media (NIM) plus SMAD inhibitors were changed every 48 hours. On day 7, once embryoid bodies began to show rosette clusters indicating early neuroepithelia morphological hallmarks, spheroids were transferred to Matrigel (Corning, 354230) coated tissue culture plates with NIM and SMAD inhibitors were removed. Media was changed every 24 hours until spheroids had sufficiently attached, and each spheroid exposed the rosette clusters within. On day 14, rosette clusters were mechanically removed and transferred to tissue culture flasks with NIM plus FGFb (Peprotech, 100-18B). Media was changed every 72 hours. On day 20, spheroids were triturated into a single cell suspension and transferred to a new untreated cell culture flask with astrocyte media (ASM) (DMEM-F12 (Gibco, 10565-018), N2 Supplement (Gibco, 17502-048), B27 -Vit.A Supplement (Gibco, 12587-010), Heparin (Stemcell Tech, 07980)) plus Y27632 (Tocris, 1254). From Day 28 to 180, spheroid aggregates were maintained in suspension with ASM plus EGF and FGFb (Peprotech, 100-15 & 100-18B) with media changes every 4-5 days. Spheroid aggregates were triturated every 7-10 days and transferred to new untreated tissue culture flasks.

### 2D and 3D cell culture

#### iNeurons Validation

iNeurons were matured in 2D monoculture for four weeks by thawing and plating previously induced, harvested, and frozen iNeurons onto PDL (Sigma, P6407-5mg) coated six-well plates. iNeuron cultures were fed with BrainPhys Complete media (BrainPhys basal (StemCell Tech, 05790), Laminin (Gibco, 23017-015), NT3 (Peprotech, 450-03), BDNF (Peprotech, 450-02), N2 (Gibco, 17502-048), B27 -Vit.A (Gibco, 12587-010), HEPES (Gibco, 15630-106)) and maintained with ~2mls/well and a 50% media change every 72 hrs.

#### iAstrocyte validation

On approximately day 120, cultures were initially validated as astrocyte progenitor cells using ICC to confirm post-mitotic behavior, stellate morphology and canonical astrocyte gene expression (sFig2 A-C). On day 0 of ICC validation, a small number of spheroids were triturated into a single cell suspension and plated on Matrigel coated glass cover-slips at ~10k cells/cm^2^ using ASM plus CNTF and BMP4 (Peprotech, 450-13 & 120-05ET). Media was changed every 48 hours. On day 7, cultures were fixed with 4% PFA and processed for ICC validation.

#### 2D co-cultures

iNeurons and iAstrocytes were plated together at 1:1 ratio on Matrigel coated 24 well plates at a collective density of ~1×10^6^ cells/well. Cells were maintained with BrainPhys Complete media at ~1ml/well with a 50% media change every 72 hrs.

#### 3D co-cultures

iNeurons and iAstrocytes were maintained, validated, and cryo-preserved in order to maximize consistency and repeatability across multiple iterations of various experiments described herein. 3D mbOrgs co-cultures were prepared by thawing or dissociating constitutive cell types in single cell suspension. Cell types were then combined at specified ratios (1:1, iAst:iN) prior to being aliquoted into round-bottom plates. Plates were briefly centrifuged and left undisturbed for ~24 hours, allowing cells to coalesce into self-assembled spheroids. Organoids were maintained with a partial media change every 72 hours using BrainPhys Complete media.

### Rescue experiment

Cells were plated as described above including an extra set of *GRN^−/−^* co-cultures which were treated with PGRN at 1μg/ml (Adipogen, AG-40A-0188Y). Media was partially changed and treatment replenished every 72 hours. Cells were then processed for RNA extraction or immunocytochemistry as described below.

### Western blot

mbOrg or 2D cultures were lysed in RIPA buffer (Thermo Fisher Scientific Cat#PI89900) complemented with Phosphatase and Protease inhibitors (Thermo Fisher Scientific Cat#1862495 and 1862209) according to manufacturer’s indications. Lysates were sonicated (QSonica, 55W, 110V Cat#Q55-110) for 10s/sample on ice and protein concentration was determined using Pierce BCA protein assay kit following manufacturer’s instruction (Thermo Fisher Scientific cat#23227). 20μg to 30 μg of total protein from each lysate was combined with loading buffer (4X, Thermo Fisher Scientific, Cat#84788) and beta mercaptoethanol (0.05 M, BioRad Laboratories Cat#1610710) and loaded into a NuPAGE 4%–12% or 10% Bis-Tris Gel (Invitrogen, Cat# NP0336BOX, NP0301BOX) alongside a protein ladder (Thermo Fisher Scientific Cat#26619) to determine protein size. The gels were run in MOPS (Thermo Fisher Scientific Cat# NP0001) or MES (Invitrogen Cat#NP0002) buffer depending on the desired protein separation. Subsequently, the gel was transferred onto a nitrocellulose membrane using Trans Blot Sd Semi-Dry Transfer Cell (BioRad Cat#1703940) according to the manufacturer’s instructions. After transfer, the membrane was stained with PonceauS, washed and blocked in 5% milk in PBS-T (PBS with 0.02% tween) or 10% BSA TBST-T (TBS + 0.02 tween), followed by overnight incubation with primary antibodies in 1% Milk or BSA at 4C. After incubation, the membrane was washed three times with PBS-T or TBS-T and then incubated with HRP conjugated secondary antibodies (Thermo Scientific Pierce Goat anti rabbit Cat#32260 or Goat anti mouse Cat#32230) at room temperature for 1hr. The membrane was then washed 3 times with PBS-T, incubated for two minutes with Pierce ECL western blotting substrate (Thermo Fisher Cat# 32106) and imaged at ChemiDoc (BioRad). Digital images were processed and analyzed using the image analysis software, ImageLab (BioRad).

**Table 1.** Primary antibodies used for western blot

| Target | Vendor and Cat# | Concentration |
| --- | --- | --- |
| GAPDH | Sigma-Aldrich, Cat# G8795 | 1:3000 |
| PGRN | ThermoFisher, cat# 40-3400 | 1:250 |
| TDP-43 | Proteintech Cat#66734-1-1g | 1:500 |
| Phospho-TDP43 (Ser409/410) | Proteintech Cat#80007-1-RR | 1:500 |
| SYN1 | Synaptic Systems Cat#101-004 | 1:300 |
| PSD95 | Abcam Cat#Ab18258 | 1:500 |

### 3D Organoid sample processing and cryosectioning for ICC

Samples were fixed in 4% PFA for ~25 minutes and washed three times with DPBS. Samples were then incubated overnight at 4°C in DPBS plus 30% sucrose and transitioned to a 1:1 solution of OCT & DPBS containing a final concentration of 30% sucrose. Samples were then incubated in OCT at RT for ~15 minutes prior to being embedded in OCT and frozen. OCT embedded samples were then cryosectioned at 20μm intervals using a Leica cryostat prior to applying the ICC protocol detailed below.

### Immunocytochemistry

For 2D cultures, cells were gently washed in DPBS and fixed in 4% PFA (EMS; 50-980-487). Samples were washed three times in DPBS and incubated with blocking buffer (Glycine 0.1M, 5% goat/donkey serum, 1% BSA, 0.25% TritonX, 100mM glycine in DPBS) for one hour at room temperature. Samples were then incubated overnight with primary antibody at the appropriate concentration in primary blocking buffer (5% goat/donkey serum, 1%BSA, 0.25% triton X-100, 100nM glycine in DPBS) at 4C overnight, washed three times in DPBS and incubated with secondary antibodies diluted in secondary blocking buffer (10% BSA in DPBS) at room temperature for 1h. Samples were washed three times in DPBS and incubated with DAPI (Sigma-Aldrich Inc Cat#D9542-5MG) for 1 min at RT and washed twice in DPBS. If on coverslips, samples were mounted on microscope slides (Fisher Scientific Cat#1255015) using ProLong Gold (Invitrogen Cat#P36934) and dried at room temperature overnight. Sections were imaged at LSM900 confocal microscope (Zeiss) or Fluorescent microscopy images were acquired using a Revolve microscope (Echo Laboratories, San Diego, CA, USA). Images were analyzed using ImageJ(FIJI) (Schindelin et al., 2012).

For 3D cultures, slices were stained as described above. Briefly, we used a hydrophobic pen to delimitate the area of staining. We then washed with PBS to clean the excess OCT and start with the protocol described above. The PGRN antibody staining protocol included an antigen retrieval treatment by incubating tissue sections in 10 mM sodium citrate (pH 6.0) at 90°C for 10 minutes before the incubation in the blocking solution. For all 3D slices, labeling was done at room temperature, but samples were treated otherwise the same as 2D cultures.

**Table 2.** Primary antibodies used for ICC

| Target | Vendor and Cat# | Concentration |
| --- | --- | --- |
| NFIA | Sigma Cat#HPA008884 | 1:200 (2D) 1:400 (3D) |
| TUJ1 | Millipore Cat#AB9354 | 1:200 |
| AQP4 | Sigma Cat#HPA014784 | 1:500 |
| CD44 | BD bioscience Cat#559046 | 1:200 |
| GFAP $\delta$ | Millipore Cat#AB9598 | 1:500 |
| TDP-43 | Proteintech Cat#66734-1-1g | 1:500 |
| Phospho-TDP43 (Ser409/410) | Proteintech Cat#22309-1-AP | 1:500 |
| PGRN | R&D Sys. Cat#AF2557 | 1:100 |
| PGRN | R&D Sys. Cat#AF2420 | 1:2000 |
| MAP2 | Abcam Cat#ab5392 | 1:500 |
| PSD 95 | Abcam Cat#ab13552 | 1:200 |
| SYP | Synaptic Sys. Cat#101004 | 1:200 |

**Table 3.** Secondary antibodies used for ICC

| Target | Vendor and Cat# | Concentration |
| --- | --- | --- |
| Goat-Anti-Mouse Alexa 568 | Thermofisher Scientific, A11004 | 1:500 |
| Goat-Anti-Chicken Alexa 647 | Thermofisher Scientific, A21449 |  |
| Goat-Anti-Rabbit Alexa 488 | Thermofisher Scientific, A11008 |  |
| Donkey-Anti-Mouse Alexa 647 | Thermofisher Scientific, A21202 |  |
| Donkey-Anti-Goat Alexa 594 | Thermofisher Scientific, A11058 |  |

### IMARIS reconstruction

IMARISx64 was used to create a 3D reconstruction of the original z-stack using the different color channels.

### Extranuclear TDP-43 analysis

TDP-43 quantifications were performed on FIJI. The DAPI channel and TDP-43 channel were each thresholded. Then, the image calculator was used to subtract the DAPI channel from the TDP-43 channel. Analyze Particles (0 μm^2^ – infinity) was used on the resulting image to quantify the number and area of the non-nuclear TDP-43. The DAPI channel was then used to create a selection that was restored on the TDP-43 channel, such that the nuclear TDP-43 was highlighted, and the intensity could be measured in that region.

### RNA isolation for bulk RNAseq and RT-PCR

RNA extraction was performed using the Trizol/phenol-Chloroform method (Sigma, T9424) as previously described (Krencik et al., 2015) and according to manufacturer specifications. Each sample contained ~50 organoids, totaling ~2.5 x10^6^ of cells per sample. The extracted RNA was used as a template for the synthesis of complementary DNA (cDNA) through reverse transcription, using iScript(tm) cDNA Synthesis Kit (Bio-Rad Cat#1708891) according to the manufacturer’s protocol.

### Quantitative PCR

cDNA samples were treated for genomic DNA contamination using TURBO DNA-free kit (Life Tech Cat#AM1907) per manufacturer’s instructions. The cDNA was then diluted to a concentration of 5 ng/l and 4 μl of each sample (total of 20 ng) were aliquoted in a white walled 96 well plate (Thermo Scientific) in technical duplicates. Samples were processed using PrimeTime std qPCR Assay (IDT) and IDT commercial standard conjugated primers for each of the genes analyzed except for cryptic STMN2 for which primers were designed by Kevin Eggan’s lab and kindly shared with us (Forward: CTCAGTGCCTTATTCAGTCTTCTC, Probe: TCAGCGTCTGCACATCCCTACAAT, Reverse: TCTTCTGCCGAGTCCCATT). The quantitative PCR was run on Bio-Rad C1000 Thermal Cycler/CF96 Real-Time System following manufacturer’s instructions. Data analysis was carried out applying the Pfaffl mathematical model for relative transcript quantification (Pfaffl, 2001) using RNA18S as a housekeeping gene.

### RNA sequencing and analysis

RNA integrity was analyzed on an Agilent 2100 Bioanalyzer using RNA 6000 Nano kit (Agilent, 5067-1511). Only samples with an RNA integrity number (RIN) ≥ 9.4 were used to perform bulk RNA sequencing. Nugen Universal Plus (Tecan) was used as a library kit and libraries were sequenced on a SP300 flow cell of the Illumina NovaSeq 6000 machine with a paired-end 150 bp sequencing strategy (average depth 90 million reads/sample) at UCSF Genomics Core Facility. Genome was aligned to Ensembl Human.GRCh38.103. Kallisto 0.46.01 was used to generate transcript abundance files for each sample. Transcript counts files for each sample were generated using txImport and transcript differential analysis was performed using DESeq2 v1.24.0. A total of 6 samples were spread across two conditions.

### Phagocytosis assay

Astrospheres were dissociated and counted. 10 wells at ~50k cells/well were plated in a 96 well/plate (~500k samples per condition). Astrocyte maturation was performed as previously described in iAstrocyte validation. One set of GRN^−/−^ astrocytes was treated with PGRN 1 μg/ml (Adipogen, AG-40A-0188Y) every 72 hours. On the day of analysis, a subset of samples per condition were pre-treated with Cytochalasin D 10μM (Sigma #C2618) for 15 minutes to inhibit actin polymerization. Phagocytosis assay was performed as previously described (Dräger et al., 2022). Briefly, all samples were incubated with pHRodoRed-labeled synaptosomes at a concentration of 1 mg/ml in astrocyte media. Synaptosomes were isolated from fresh Innovative Grade US Origin Rat Sprague Dawley Brain (Innovative Research, Inc.; Cat. No. IGRTSDBR) with the Syn-PER™ Synaptic Protein Extraction Reagent (Thermo Scientific™; Cat. No. 87793) according to the manufacturer’s protocol and then labelled with pHrodo™ Red, succinimidyl ester (pHrodo™ Red, SE) (ThermoFisher Scientific; Cat. No. P36600) as previously described (Dräger et al., 2022). Cells were washed twice with DPBS, dissociated, resuspended in ice-cold DPBS, and analyzed via flow cytometry. Flow cytometry data were analyzed using FlowJo (https://www.flowjo.com/).

### Statistical analysis

Statistical analyses were done using Prism 9.0 (GraphPad). If normally distributed, two-tailed unpaired Student’s *t*-test was performed. If normally distributed, One way Anova test was performed. *P*<0.05 was considered significant. If non-normally distributed, Mann-Whitney *U*-test was performed. *P*<0.05 was considered significant.

Using DESeq2, Wald tests were performed to evaluate genes for differential expression between conditions. FDR multiple testing correction method was then used for adjusted p-values.

## Acknowledgments

We thank Arnab Ghosh and Efy Hernandez of the UCSF Genomics CoLab for the RNA-seq Library preparation and raw analysis. We would also like to thank Su Ling Wang for her contribution on the figure graphics. We also thank Bruce Conklin for support and advice.

## Autor Contributions

M.dM., M.Koontz, and E.M.U. designed experiments. M.dM., M.Koontz, E.M., N.S., A.R., Y.K., S.L.G., N.M.D. and K.L. performed experiments and/or analyzed data. Y.M. generated the GRN^−^/^−^ iPSC lines. M.dM, E.M.U., E.M., H.L., K.S. conceived the hypothesis. E.M., J.R.K, M.Kurnellas, M.Kampmann, M.E.W., E.J.H and E.M.U. provided resources. M.dM., M.Koontz, E.M. and E.M.U. wrote the manuscript. All authors reviewed and approved the manuscript.

