## Supplementary Figure for "Granulin Loss of Function in Human Mature Brain Organoids Implicates Astrocytes in TDP-43 Pathology"

de Majo, Koontz, et al.

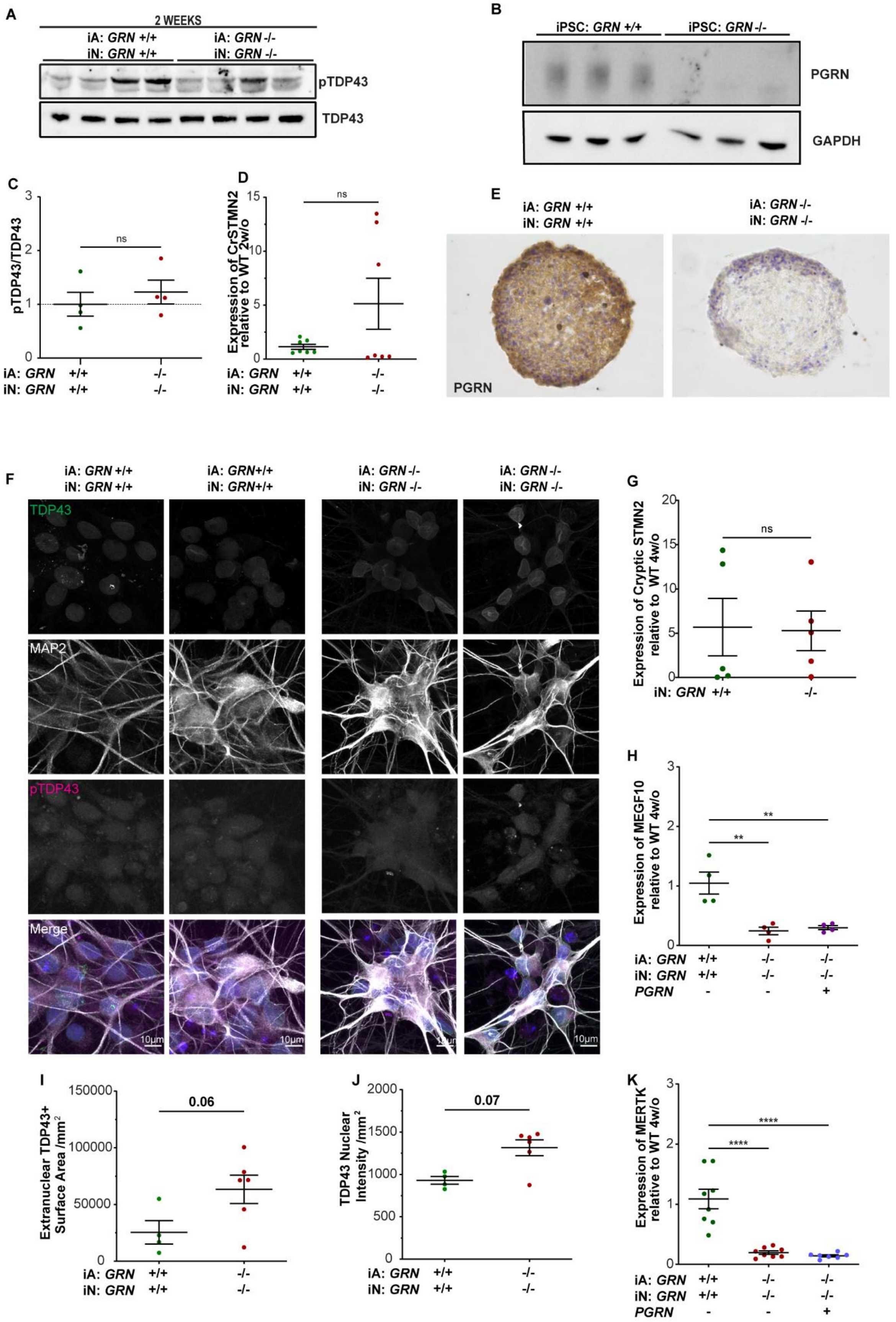

A
2 WEEKS
iA: GRN +/+
iN: GRN +/+
iA: GRN -/-
iN: GRN -/-
pTDP43
TDP43
B
iPSC: GRN +/+
iPSC: GRN -/-
PGRN
GAPDH
C
pTDP43/TDP43
ns
D
Expression of CrSTMN2 relative to WT 2w/o
ns
E
iA: GRN +/+
iN: GRN +/+
iA: GRN -/-
iN: GRN -/-
PGRN
F
TDP43
MAP2
pTDP43
Merge
10µm
G
Expression of Cryptic STMN2 relative to WT 4w/o
ns
iN: GRN +/+ -/-
H
Expression of MEGF10 relative to WT 4w/o
**
**
PGRN
I
Extranuclear TDP43+ Surface Area /mm2
0.06
J
TDP43 Nuclear Intensity /mm2
0.07
K
Expression of MERTK relative to WT 4w/o
****
****

**Supplementary figure 1.** A) Western blot of mbOrg whole lysate showing similar expression on pTDP-43 in *GRN*$^{-/-}$ and *GRN*$^{+/+}$ mbOrg when normalized to total TDP-43 at two-week timepoint. C) Western blot quantification showing no difference in expression on pTDP-43 in *GRN*$^{-/-}$ compared to *GRN*$^{+/+}$ mbOrgs when normalized to total TDP-43 (n=4, unpaired t test, two tailed, p=ns). B) Western blot of WTC11 iPSCs whole lysate confirming GRN knock out in GRN$^{-/-}$ WTC11 lines. D) Quantification of CrSTMN2 expression using qPCR showing no difference in CrSTMN2 expression in *GRN*$^{-/-}$ compared to *GRN*$^{+/+}$ mbOrg after two weeks in culture. E) Brightfield images of PGRN immunostaining in 3D cultures at 2-week timepoint showing the loss of PGRN in GRN$^{-/-}$ pure combination compared to *GRN*$^{+/+}$ pure combination. F) Representative ICC images of 2D neuronal cultures after for weeks. Cells were stained for TDP-43, MAP2 and pTDP-43 (scale bar 10µm). G) Quantification of CrSTMN2 expression using qPCR showing no difference in CrSTMN2 expression in GRN$^{-/-}$ compared to *GRN*$^{+/+}$ 2D neuronal cultures after four weeks in culture. H and K) Quantification of MERTK and MEGF10 expression using qPCR showing significantly MERTK and MEGF10 lower expression in *GRN*$^{-/-}$ compared to *GRN*$^{+/+}$ 2D cocultures after four weeks in culture. MERTK and MEGF10 expression is not rescued when *GRN*$^{-/-}$ co-cultures were treated with exogenous PGRN. I and J) quantification of extranuclear TDP-43 per mm$^2$ (I) and TDP-43 nuclear intensity per mm$^2$ (J) in *GRN*$^{-/-}$ compared to *GRN*$^{+/+}$ mbOrg. Each dot represents one independent mbOrg (*GRN*$^{+/+}$ n=4, *GRN*$^{-/-}$ n=6, unpaired t test, two tailed, $p<0.05$). For all graphs data are represented as mean ± SEM.

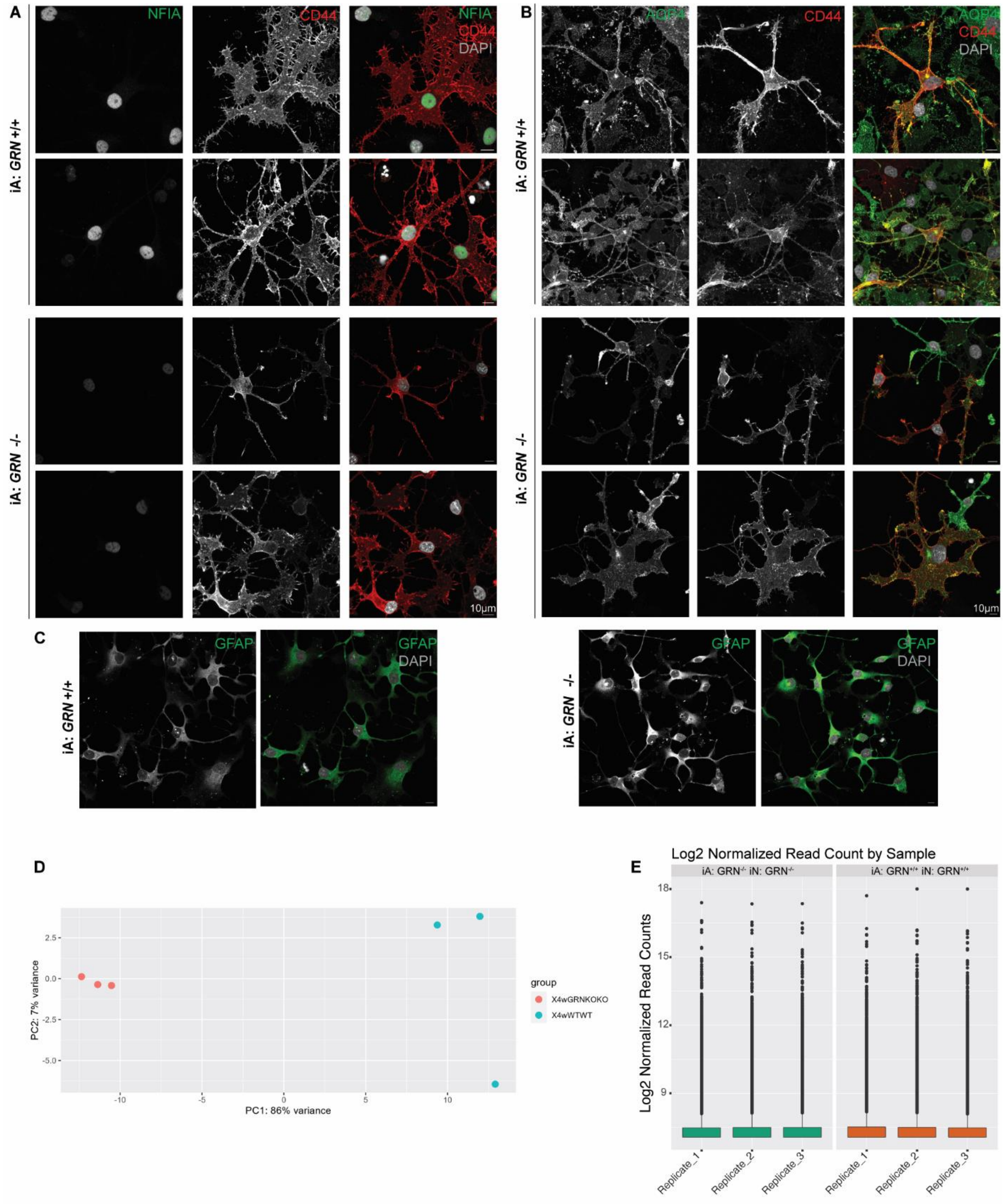


**F**

| | | |
|---|---|---|
| *GRN +/+* | Parental | cccgccagGCTGTGTGCT GCGAGGATCGC CAGCACTGCT GCCCGGCTGG CTACACCTGC AACGTGAAG-------GCTCGATCCTGCGA GAA |
|  | Parental | cccgccagGCTGTGTGCT GCGAGGATCGC CAGCACTGCT GCCCGGCTGG CTACACCTGC AACGTGAAG-------GCTCGATCCTGCGA GAA |
| | | *Exon 12* |
| *GRN -/-* | Allele 1 | cccgccagGCTGTGTGCT GCGAGGATCGC CAGCACTGCT GCCCGGCTGG CTACACCTGC AACGTGAAG-------GCTC---------- GAA (10 bp deletion) |
| | Allele 2 | cccgccagGCTGTGTGCT GCGAGGATCGC CAGCACTGCT GCCCGGCTGG CTACACCTGC AACGTGAAGGCTCGATGCTCGATCCTGCTA GAA (7 bp insertion; 1 bp mutation) |

**Supplementary figure 2.** A) Representative ICC images of 2D iA cultures stained for NFIA, CD44 and DAPI (scale bar 10µm) after one week differentiation B) Representative ICC images of 2D iA cultures stained for AQP4, CD44 and DAPI (scale bar 10µm) after 1 week differentiation. C) Representative ICC images of 2D iA cultures stained for GFAP and DAPI (scale bar 10µm) after one week differentiation. D) Principal components plot indicating distance between samples with principal component 1 (PC1) vs principal component 2 (PC2) and highlighting samples separation by genotype. E) Boxplots by sample for normalized read count values for all six analyzed samples. F) Diagram showing the *GRN* region in the WTC11iPSC line that was modified in the $GRN^{-/-}$ WTC11 iPSC. The modification results in a 10 base pair deletion (allele 1) and a 7+1 base pair insertion (allele 2) in Exon 12. These nonsense variants lead to the formation of a premature stop codon and a transcript that is degraded by nonsense mediated decay resulting in a *GRN* knock out iPSC line. For all graphs data are represented as mean ± SEM.
